# Heat shock induces the depletion of Oct4 in mouse blastocysts and stem cells

**DOI:** 10.1101/264044

**Authors:** Mo-bin Cheng, Xue Wang, Yue Huang, Ye Zhang

**Affiliations:** State Key Laboratory of Medical Molecular Biology, Chinese Academy of Medical Sciences & Peking Union Medical College, Beijing 100005, China; Department of Biochemistry and Molecular Biology, Institute of Basic Medical Sciences, Chinese Academy of Medical Sciences & Peking Union Medical College, Beijing 100005, China; Department of Medical Genetics, Institute of Basic Medical Sciences, Chinese Academy of Medical Sciences & Peking Union Medical College, Beijing 100005, China

**Keywords:** Oct4, heat, shock, Dapk1, Pin1

## Abstract

Temperature is an important microenvironmental factor that functions epigenetically in normal embryonic development. However, the effect of heat shock in the stem cells is not fully understood. Oct4 is a tightly regulated master regulator of pluripotency maintenance in stem cells and during early embryonic development. We report here that Oct4 protein level was significantly reduced under heat shock in mouse blastocysts and embryonic stem cells. The reduction in Oct4 in the mouse embryonic stem cells under heat shock was mediated by a ubiquitin-proteasome pathway that was dependent on the activity of death-associated protein kinase 1 (Dapk1) to phosphorylate its substrate, Pin1. Our results imply that the depletion of Oct4 via brief heat shock, such as a high fever, during early pregnancy might severely impair the growth of the mammalian embryo or even cause its death.

## Introduction

Oct4 is a master transcriptional regulator that maintains the naive pluripotency of stem cells. A deficiency in Oct4 fails to form a pluripotent inner cell mass (ICM) that results in the pre-implantation lethality of the mouse embryo (Nichols et al, 1998). Oct4 is capable of reprogramming various somatic cell types into pluripotent cells, either in combination with other transcription factors (Takahashi & Yamanaka, 2006) or on its own (Kim et al, 2009). The maintenance of an appropriate level of Oct4 in stem cells leads to either the maintenance of pluripotency or cell lineage differentiation (Lu et al, 2009; Niwa et al, 2000; Plachta et al, 2011; Radzisheuskaya et al, 2013).

An appropriate temperature is known to be a micro-environmental or epigenetic factor in normal embryonic development in the animal kingdom (Coffey et al, 2006); in contrast, heat shock is an established teratogen in mammals, including humans (Graham et al, 1998). However, we know next to nothing regarding the effect and mechanism of heat shock on stem cells or embryonic development.

Here, we report hyperthemia induces depletion of Oct4 in mouse blastocysts and stem cells, which is mediated by a ubiquitin-proteasome pathway that is dependent on heat shock activated Dapk1/Pin1 pathway.

## Results And Discussion

### Heat shock depletes Oct4 in mouse blastocysts and stem cells

Because the maintenance of a pluripotent ICM is Oct4 dependent and the ICM is more sensitive to elevated temperatures in the blastocyst than in the trophectoderm (Amano et al, 2000), we thus determined the expression levels of the pluripotency-associated transcription factors such as Oct4 in the ICM after heat shock. Mouse blastocysts were pretreated at 39 or 42°C for 1 hour, and the expression levels of the transcription factors were then detected by immunofluorescence staining. We showed that although the Oct4 level in the ICM diminished significantly after 42°C treatment, Klf4 and Sox2 remained largely unchanged. Nanog was reduced compared with the control after treatment at 37°C (Fig. 1A). These results indicated that heat shock produces the largest significant decrease of Oct4 expression during early embryonic development.

**Fig. 1.**
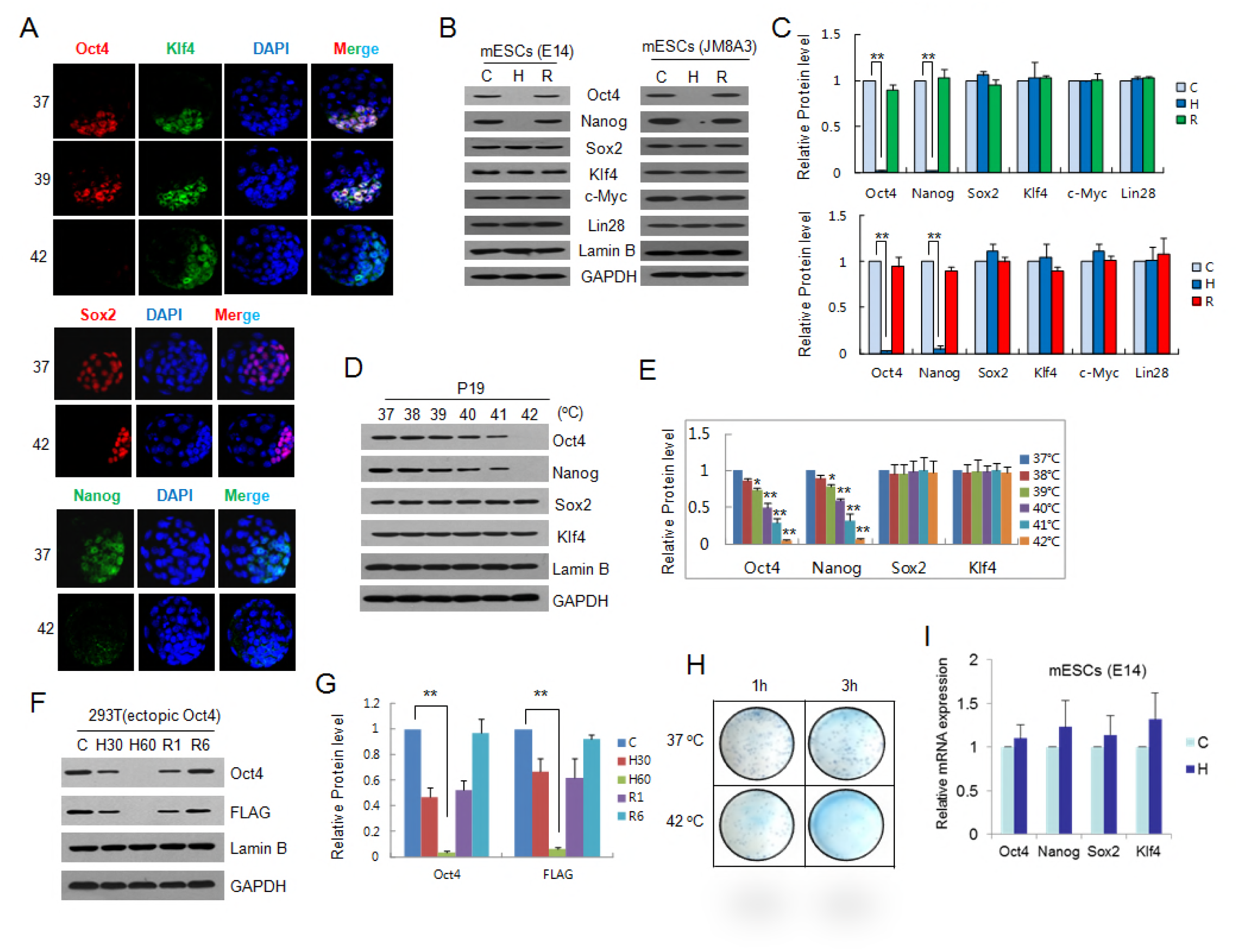
Impact of heat shock on Oct4 expression in mouse blastocysts and ESCs. **(A)** Immunostaining of Oct4, Klf4, Sox2 and Nanog in the ICMs of E3.5 blastocysts. The blastocysts were incubated for 1 h at the indicated temperature, and the nuclei were shown by DAPI staining (blue). **(B and C)** Western blot analysis of Oct4, Nanog, Sox2, Klf4, c-Myc, and Lin28 from E14 and JM83A mESCs. Lamin B and GAPDH were used as controls. C: 37 °C; H: 42 °C for 1 h; R: recovery at 37 °C for 6 h following 42 °C treatment (B). Quantification data represent the mean ± SD of three independent experiments. ∗p < 0.05, ∗∗p < 0.01 (C). **(D and E)** Western blot analysis of Oct4, Nanog, Sox2, and Klf4 from P19 cells incubated for 1 h at each indicated temperature (D). Quantification data represent the mean ± SD of three independent experiments. ∗p < 0.05, ∗∗p < 0.01 (E). **(F and G)** Western blot analysis of ectopically expressed FLAG-tagged Oct4 from HEK293 T cells (F). Quantification data represent the mean ± SD of three independent experiments. ∗p < 0.05, ∗∗p < 0.01 (G). **(H)** Colony formation assays of JM8A3 mESCs treated as indicated. The colonies were visualized using methylene blue staining. **(I)** Heat shock produced no obvious effects on the mRNA expression of oct4, nanog, sox2, or klf4 in E14 mESCs, as revealed by real-time RT-PCR assays. Each bar represents the mean value obtained from at least three independent experiments normalized against gapdh mRNA; the S.D. is shown on top of each bar.

Next, we investigated the effects of heat shock on the expression of the key pluripotency transcription factors in embryonic stem cells (ESCs). We showed in mouse ESCs (E14) that the Oct4 protein expression was remarkably reduced under heat shock (42°C for 1 hour), and western blotting showed a full recovery 6 hours after the removal of the cells from the stress. Similar results were demonstrated for Nanog, albeit to a lesser extent (Fig. 1B, left panel and Fig. 1C, upper panel). A similar profile was also shown in the JM83A mouse ESCs (Fig. 1B, right panel and Fig. 1C, lower panel). In contrast to Oct4 and Nanog, other important pluripotency factors, such as Sox2, Klf4, c-Myc, and Lin 28, were not affected at the protein level by heat shock. We then treated P19 cells (mouse embryonic carcinoma cells) with a series of temperatures from 38°C to 42°C to verify that an Oct4 reduction could be caused by temperatures elevated to the fever range. The results demonstrated that Oct4 and Nanog were reduced in the P19 cells only at 40°C or higher (Fig. 1D and 1E). HEK293T cells transfected with FLAG-tagged Oct4 were further subjected to heat treatment, and the thermal-dependent reduction of ectopic FLAG-tagged Oct4 was clearly identified by antibodies against either Oct4 or its FLAG-tag (Fig. 1F and 1G). To assess the impact of heat shock on the self-renewal of mouse ESCs (mESCs), a colony formation assay was performed. It was shown that colony formation was severely impaired by heat shock at 42°C (Fig. 1H). We thus suggest that the heat shock-mediated reduction in Oct4 is associated with the inhibition of self-renewal in mESCs.

### Thermal degradation of Oct4 is Dapk1/Pin1 pathway dependent

To determine how the Oct4 reduction is regulated under heat shock, we first demonstrated that the mRNA levels of *Oct4, Nanog, Sox2 and Klf4* in E14 mESCs were not obviously affected under heat shock using real-time RT-PCR (Fig. 1I) and RNA-Seq (data not shown). In contrast, the proteasome inhibitor MG132 could prevent the Oct4 reduction under heat shock in E14 mESCs (Fig. 2A), suggesting that the rapid degradation of Oct4 is proteasome dependent. In addition, the presence of MG132 significantly induced the accumulation of polyubiquitinated Oct4 under heat shock in mESCs (Fig. 2B). We thus confirm that the degradation of Oct4 is mediated by a ubiquitin-proteasome pathway.

**Fig. 2.**
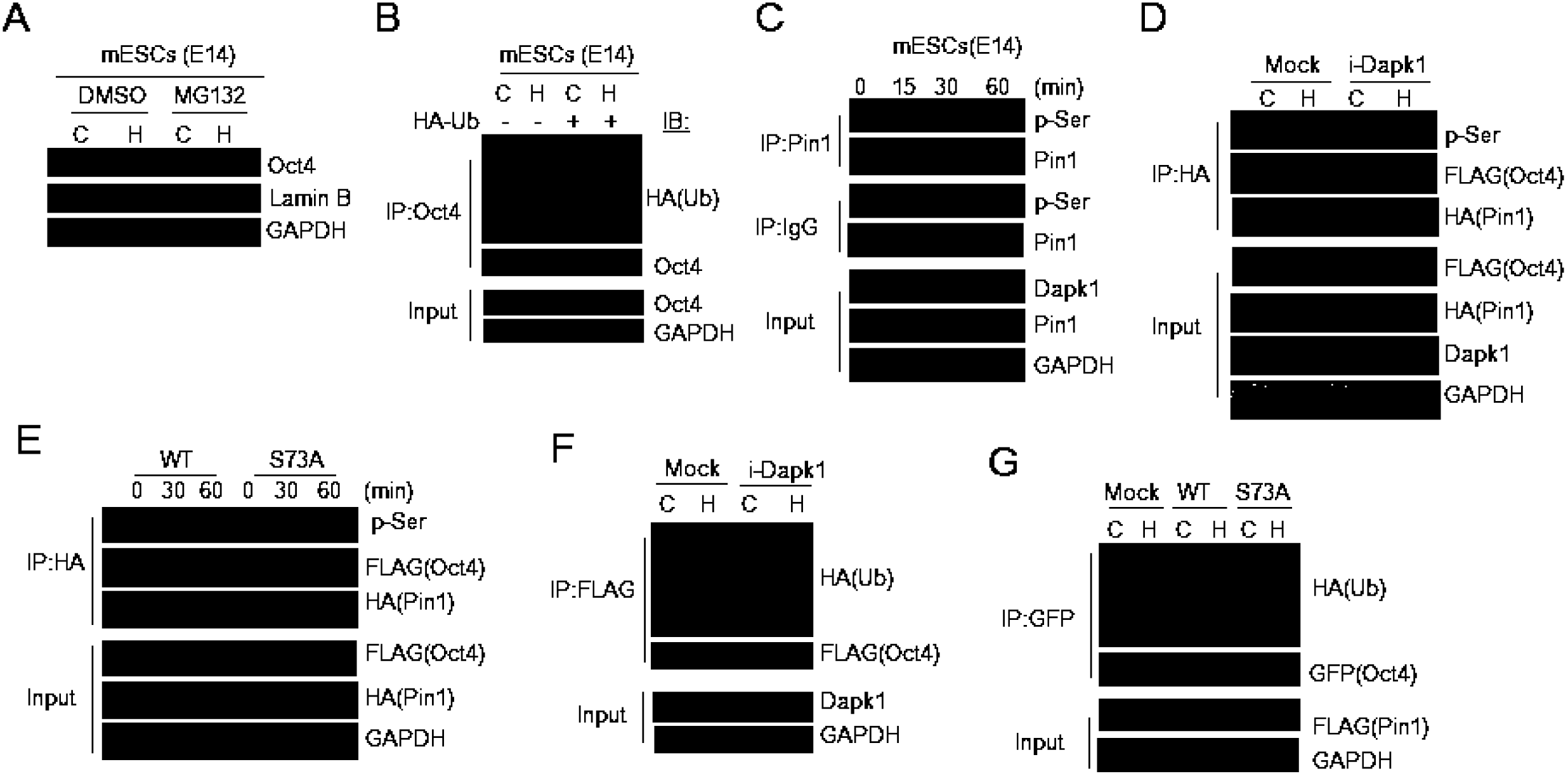
The heat shock-promoted degradation of Oct4 mediated by Dapk1/Pin1 dependent ubiquitin-proteasome pathway. **(A)** MG132 prevented the degradation of Oct4. **(B)** Heat shock induced the accumulation of ubiquitinated Oct4 in the presence of MG132. Whole cell extracts (WCEs) were immunoprecipitated (IP) with anti-Oct4 and were blotted (IB) with anti-HA and anti-Oct4. **(C)** Heat shock facilitated the phosphorylation of Pin1 in E14 mESCs. IP extracts of Pin1 and IgG (negative control) were blotted with antibodies against pan-phosphorylated serine (p-Ser) and Pin1. The inputs were blotted with antibodies against Dapk1, Pin1, and GAPDH. **(D)** Knockdown of Dapk1 abrogated both the phosphorylation of Pin1 and the degradation of Oct4 in HEK293 T cells ectopically expressing FLAG- Oct4 and HA-Pin1. WCEs were immunoprecipitated with anti-HA and blotted with the antibodies indicated on the right. **(E)** A Pin1-S73A mutant blocked the heat shock-mediated degradation of Oct4. HEK293 T cells transfected with FLAG-Oct4 and either HA-tagged Pin1 (WT) or mutant Pin1 (S73A) were treated for the indicated minutes (min). **(F)** Knockdown of Dapk1 prevented the ubiquitination of Oct4 in cells ectopically expressing FLAG-Oct4 and HA-Ub. **(G)** A Pin1-S73A mutant prevented the ubiquitination of Oct4 in HEK293 T cells ectopically expressing GFP-Oct4 and HA-Ub.

To explore the mechanism of the thermal degradation of Oct4, the death associated protein kinase 1 (Dapk1)/Pin1 pathway was determined. Pin1 is a prolyl cis/trans isomerase that functions to regulate the stability of its substrates via phosphorylation-dependent ubiquitination mediated degradation (Liou et al, 2011). Pin1 is indispensable for the self-renewal and maintenance of pluripotent stem cells (Moretto-Zita et al, 2010; Nishi et al, 2011). Dapk1 phosphorylates human Pin1 at Ser71 (mouse Ser73), which is a catalytic site that inactivates Pin1 isomerase activity (Lee et al, 2011). We demonstrated that during heat shock, the level of phosphorylated Pin1 gradually increased, which is in accordance with the accumulation of Dapk1, whereas there was no obvious change in the level of Pin1 in E14 mESCs (Fig. 2C). In contrast, the heat shock-induced phosphorylation of Pin1 was prevented by the knockdown of Dapk1, which protected Oct4 from degradation. However, the interaction between Pin1 and Oct4 remained unchanged (Fig. 2D). We next cloned an expression construct of mouse Pin1 with an S73A mutation and found that the non-effective S73 phosphorylation site in the Pin1 mutant could again prevent Oct4 degradation under heat shock (Fig. 2E). We also showed that either the knock down of Dapk1 (i-Dapk1) or the mutation of the Pin1 phosphorylation site abolished the accumulation of polyubiquitinated ectopic Oct4 (Fig. 2F and 2G). These results indicated that phosphorylation interfered with the prolyl isomerase activity of Pin1 on stabilizing Oct4, which was thus degraded via a ubiquitin/proteasome pathway. We suggested that Dapk1 abrogate stem cell maintenance and embryonic development via its phosphorylation of the Pin1-S73, which promote the degradation of Oct4 under heat shock.

To explore whether DAPK1 regulates Oct4 expression in heat shock condition during embryonic development, we detected the expression of Oct4 in *Dapk1*^-/-^ blastocysts upon heat shock at 42°C for 1 h. We collected *Dapk1*^-/-^ embryos via a two-steps procedure as described in Experimental Procedures. To distinguish the DAPK1-null embryos from the others with different genotypes by using DAPK1 immunofluorescence, we tried several times but failed, which most likely due to lack of enough blastocysts.

In fact, heat shock has been used for cancer therapy for more than a century. Numerous clinical and basic studies have shown that heat shock can significantly alter tumor cell survival (Dewhirst et al, 2005).

However, we provide the first evidence on the changes in stem cells under heat shock conditions, which suggest, similar to a high fever, heat shock could severely impair the growth of a mammalian embryo in early pregnancy or even cause its death.

Whether the reduction in Nanog in certain cell types is secondary to the depletion of Oct4 under heat shock or whether Nanog is degraded via a similar or alternate mechanism is an interesting topic remained to be explored.

In summary, heat shock could promote the degradation of Oct4 in mouse ICM, ESCs, and P19 cells. S73- phosphorylation blocked the prolyl cis/trans isomerase activity of Pin1, thereby destabilizing Oct4 for ubiquitin-proteasome-mediated degradation. These heat shock-induced changes impaired the renewal of stem cells during early embryogenesis.

## Materials and Methods

### Cell lines and cell culture

Mouse E14 ESCs were cultured in KnockOut DMEM (Life Technologies, Grand Island, NY) supplemented with 15% FCS (Sigma, St. Louis, MO), 1× GlutaMAX (Gibco, Grand Island, NY), 1× PEN/STREP (Millipore, Philadelphia, PA), 20 μM β-mercaptoethanol (Millipore), 1% nonessential amino acids (Millipore), 1% nucleotide mix (Millipore), and 100 U/ml Leukemia inhibitory factor (LIF, Millipore) on gelatinized tissue culture plates at a density of 2.5-5×10^4^/cm^2^. Mouse JM8A3 ESCs (kindly provided by Dr. Allan Bradley) were cultured in a maintenance medium consisting of KnockOut DMEM supplemented with 15% KnockOut Serum Replacer (Life Technologies, 0.1 mM β-mercaptoethanol, 0.1 mM nonessential amino acids, 1 mM glutamine, and 1000 U/ml LIF. HEK293T cells were maintained in RPMI 1640 supplemented with 10% fetal calf serum. P19 embryonic carcinoma cells were maintained in α-MEM supplemented with 10% fetal calf serum.

### Antibodies and plasmids

Antibodies against Oct4, Lamin B and GAPDH were purchased from Santa Cruz Biotech (Santa Cruz, CA). Sox2, c-Myc, Pin1 and Dapk1 antibodies were purchased from Abgent Biotech (San Diego, CA). Klf4 and Lin28 antibodies were purchased from Cell Signaling Technology, Inc. (Danvers, MA). FLAG FLAG-tagged and HA HA-tagged antibodies were purchased from Sigma.

The FLAG-tagged Pin1 and HA-tagged Pin1 eukaryotic expression plasmids were constructed by cloning Pin1 CDS into pCDNA6-FLAG and pCDNA6-HA vectors, respectively, using a PCR product amplified from a mouse ESC cDNA library. Specific primers (forward, GGGGTACCATGGCGGACGAGGAGAAG; reverse, CGGGATCCTCATTCTGTGCGCAGGAT) were used for the PCR amplification. We created a point mutation in amino acid 73 (S to A) in full-length FLAG-Pin1 or HA-Pin1. Specific primers (forward, TCACGGCGGCCCGCGTCCTGGCGGCAG; reverse, CTGCCGCCAGGACGCGGGCCGCCGTGA) were used for the PCR amplification. The FLAG/GFP-tagged Oct4 eukaryotic expression plasmid was a gift from Dr. Wenji Dong of Peking Union Medical College. The HA-Ubiquitin (Ub) eukaryotic expression plasmid was a gift from Dr. Linfang Wang of Peking Union Medical College. The oligonucleotide sequences of the Dapk1 small hairpin RNA (shRNA) were designed using Block-iT RNAi Designer (GGACACCTCCATTACTCATTGT, Life Technologies) and were connected with a HindIII/BamHI pRS vector.

### Mouse blastocyst collection and immunofluorescence

Female mice (C67BL/6N) were superovulated by injections of pregnant mare serum gonadotropin and human chorionic gonadotropin. The superovulated females were mated with male mice (B6D2F1), and the E3.5 blastocysts were flushed out of the uterus.

The immunofluorescence analysis of the blastocysts was performed as previously described (Miyanari & Torres-Padilla, 2012), with minor modifications. Briefly, the blastocysts were placed in a drop of M16 medium under mineral oil and were incubated at 37°C, 39°C and 42°C in a humidified atmosphere containing 5% CO_2_. The blastocysts were fixed with 4% paraformaldehyde for 20 min at room temperature (RT). After being washed with PBS, the blastocysts were permeabilized with 0.5% Triton X- 100 for 10 min and were then incubated in blocking solution (0.2% BSA) for 30 min. The blastocysts were incubated with a primary antibody overnight at 4°C, washed three times with 0.01% Triton X-100 and then incubated in a blocking solution containing a secondary antibody for 1 h at RT. After being washed three times with 0.01% Triton X-100, the blastocysts were mounted on glass slides with DAPI. Images were obtained using an OLYMPUS FV1000 confocal microscope. Antibodies were used for immunofluorescence of Nanog (AB5731, Millipore) and Klf4 (sc-20691, Santa Cruz).

We collected *Dapk1*^-/-^ embryos via a two-steps procedure: 1) generated *Dapk1*^+/-^ mice by crossing *Dapk1*^flox/+^ mice with *EIIa-cre* transgenic mice [strain name: FVB/N-Tg (EIIa-cre) C5379Lmgd/J); *Dapk1* ^flox/+^ mice, in which the exon3 of one *Dapk1* allele was flanked by two loxP sites, were kindly provided by Prof. Youming Lu (Huazhong University of Science and Technology, Wuhan)]. 2) Female *Dapk1*^+/-^ mice were superovulated by treating them with pregnant mare serum gonadotropin and human chorionic gonadotropin, then were crossed with male *Dapk1*^+/-^ mice. The blastocysts were flushed out of the uterus at E3.5.

### Immunoprecipitation (IP) and immunoblot analyses

Co-IP analyses were performed with ∼500 μg of proteins incubated with specific antibodies for 2 h at 4°C. Next, 20 ml of Protein A (or G)-agarose was added and incubated at 4°C overnight. The pellets were then washed with RIPA buffer, 40 μl of 1× Laemmli buffer was added, and then they were suspended and boiled. The samples were separated by SDS-PAGE and were sequentially analyzed by western blotting with individual antibodies. For quantification of Western blots, the membranes were scanned for analysis of the relative level of protein expression. The intensity of each protein band was determined by densitometry and divided by GAPDH gray value for normalization.

### Colony formation assays

The JM83A ESCs were suspended in culture medium after trypsinization and treated with a 37°C or 42°C water bath for 1, 3 or 6 h. Subsequently, and then seeded 1000 cells on one well of six-well plate (100 cells/cm^2^). After 8 d culture, the ESC colonies were stained with methylene blue (1% in 75% ethanol) for 15 min, followed with washing with 75% ethanol for 1 min.

### Quantitative real-time RT-PCR (RT-qPCR)

RT-qPCR was performed as previously described (Li et al, 2007; Wu et al, 2009). The relative expression levels of the *Oct4, Nanog, Sox2, and Klf4* genes were normalized against GAPDH using the comparative CT method, according to the manufacturer's instructions (Roter-Gene RG-3000A Real-Time PCR System, Corbett Research, Australia). The experiments were repeated at least three times. All the data from the experiments are expressed as the means ± SD.

### Ubiquitination assay

The mESCs were transfected with combinations of plasmids, including HA-ubiquitination (HA-Ub). After incubation for 42 h, the cells were treated with MG132 (25 mM) for 6 h. Whole cell extracts (WCEs) were immunoprecipitated with an antibody against Oct4 and were blotted with an antibody against HA. The HEK293T cells were co-transfected with combinations of plasmids, including HA- ubiquitination (HA-Ub) and FLAG/GFP-Oct4. After incubation for 42 h, the cells were treated with MG132 (25 mM) for 6 h. The WCEs were immunoprecipitated with an antibody against FLAG or GFP and were blotted with an antibody against HA.

### Statistical Analysis

Statistical analysis was performed using two-tailed Student's t-test. All data are shown as the mean with standard deviations (SD) from at least three independent experiments. Probabilities of P ≤ 0.05were considered as significant (*) and P ≤ 0.01 as highly significant (**).

## Acknowledgements

We thank Dr. W Dong, and L Wang for kindly providing plasmids. We thank Dr. Y Lu (Huazhong University of Science and Technology, Wuhan) for kindly providing *Dapk1* ^flox/+^ mice. This work was supported by the CAMS Initiative for Innovative Medicine (2016-I2M-3-002 to YZ and YH) and the National Natural Science Foundation of China (31371305 to YZ; 31021091 to YH).

## Author contributions

Y.H. and Y.Z. designed the studies, conducted the experiments, interpreted the data and wrote the manuscript. M.C. conducted the biochemical studies. X.W. conducted the blastocyst studies.

## Conflict of Interest

The authors declare that they have no conflict of interest.

